# Widespread distribution of the BpfA-carrying bacteria involved in the bisphenol F degradation in Pseudomonadota and Actinomycetota

**DOI:** 10.1101/2025.06.02.657393

**Authors:** Changchang Wang, Mingliang Zhang, Yanni Huang, Qian Li, Junqiang Hu, Kaihua Pan, Qian Zhu, Wankui Jiang, Jiguo Qiu, Xin Yan, Qing Hong

## Abstract

The global use of bisphenol F (BPF) has led to widespread environmental contamination, posing significant threats to ecosystems and human health. However, the genes involved in BPF degradation remained unknown. In this study, a novel oxidase gene, *bpfA*, was obtained from strain *Microbacterium* sp. F2 through a four-step purification strategy. BpfA is classified as a member of the 4-phenol oxidizing (4PO) subfamily and contains a conserved FAD-binding domain (Asp152 and His393) as well as a Tyr-Tyr-Arg triad (Tyr93, Tyr474, Arg475). BpfA catalyzes the conversion of BPF to 4,4’-dihydroxydibenzophenone (DHBP) through a series of three consecutive reactions. In addition, BpfA exhibits catalytic activity towards 4-alkylphenols, such as vanillyl alcohol (VA), 4-n-propylguaiacol (4PG), and 4-(methoxymethyl)phenol (4MOP). BpfA homologs are widely distributed in the environment, particularly in soil. Bioinformatics-based taxonomic profiling revealed that BpfA homologs are widely distributed in metagenomes from cultivated land and forests, mainly belonging to Pseudomonadota and Actinomycetota. This study enhances our understanding of the microbial degradation mechanism of BPF, elucidates the catalytic function of the novel oxidase gene *bpfA* and its distribution pattern in the environment, and provides important insights into the evolutionary origin of BPF degradation genes and the targeted mining of environmental microbial resources.

## Introduction

Bisphenols (BPs) are one of the important chemical raw materials for the synthesis of high-molecular-weight materials and are widely used in various consumer products, including bisphenol A (BPA), bisphenol F (BPF), bisphenol B (BPB), bisphenol S (BPS), and bisphenol AF (BPAF). However, toxicological studies have shown that BPA has endocrine-disrupting effects and may trigger a variety of health problems, such as cancer [1], reproductive disorders [2,3], and obesity [4,5]. As one of the main substitutes for BPA, the usage of BPF has increased year by year, and it has become a new, widely-present pollutant in water bodies [6]. BPF enters aquatic, soil, and atmospheric environments via industrial effluent discharge, plastic product degradation, and landfill leachate and has been widely detected globally [7,8]. BPF contamination in East Asian aquatic systems reaches notable levels, exceeding 1000 ng/L in rivers and seawater across Japan, Korea, and China; the highest concentration (2850 ng/L) was detected in Tokyo’s Tamagawa River [9–11]. Moreover, the breast milk samples collected from 181 healthy women across various provinces and cities in China revealed an average BPF concentration of 0.107 μg/L [12]. This suggests that BPF, in addition to its widespread presence in the aquatic environment, may also pose risks to human health through bioaccumulation.

BPF has toxic effects on organisms. Exposure to 0.5 mM BPF during the embryonic stage of fruit flies reduced the number of neuroblasts and intermediate neural progenitor cells, leading to an imbalance in the neuron-to-glial cell ratio. This disrupted the proliferation of neural stem cells, resulting in neurodevelopmental toxicity and abnormal larval behavior [13]. Through transcriptomic and metabolomic studies, it was found that exposure to 1.0 μM BPF upregulated the expression of pro-inflammatory cytokines (such as IL-17A and TNF-α) in human colonic epithelial cells (NCM460), while downregulating tight junction proteins (ZO-1 and CLDN1), thereby disrupting intestinal barrier function [14]. In addition, adult male zebrafish exposed to BPF concentrations of 500 nM and 2500 nM exhibited significant reductions in sperm count, decreased levels of serum testosterone (T), and increased levels of vitellogenin (VTG) in the liver, indicating that BPF has endocrine-disrupting effects on adult male zebrafish [15].

Several bacterial strains capable of growing with BPF have recently been identified, including *Pseudomonas* sp. HS-2 [16], *Pseudomonas* sp. ZH-FAD [17], *Sphingobium yanoikuyae* FM-2 [18], *Sphingobium yanoikuyae* TYF-1 [19], *Arthrobacter* sp. YC-RL1 [20], and *Microbacterium* sp. F2 [21]. The identification of similar intermediates in the BPF degradation process among strains from various genera has led to the proposal of a common catabolic pathway (Fig. S1). This pathway involves the initial hydroxylation of the BPF carbon bridge to form (4-hydroxyphenyl)methanol (BHPM), followed by dehydrogenation to produce 4,4 ’ - dihydroxydibenzophenone (DHBP). Subsequently, the Baeyer-Villiger reaction occurs between the two phenolic rings to generate 4-hydroxybenzoic acid-4-hydroxyphenyl ester (HPHB), which is hydrolyzed into *p*-hydroxybenzoic acid (PHBA) and hydroquinone (HQ). These compounds are further degraded to provide carbon sources for the growth of the strains. Although several BPF-degrading bacterial strains have been reported, the specific genes involved in the conserved BPF metabolic pathway remain unknown. Elucidating these BPF degradation genes is crucial for understanding the transformation and ultimate fate of BPF in the environment.

In the current study, a novel oxidase gene, *bpfA*, was identified in the previously isolated BPF-degrading strain *Microbacterium* sp. F2, which catalyzes the conversion of BPF to DHBP through three consecutive reactions. The classification of the *bpfA* gene and its distribution in various environments worldwide were analyzed. These results provide novel perspectives on the variety of BPF degradation genes across distinct microbial strains.

## Materials and methods

### Chemicals and media

Bisphenol F (BPF), bisphenol E (BPE), vanillyl alcohol (VA), 4,4’-dihydroxybenzophenone (DHBP), 4-*n*-propylguaiacol (4PG), 4-hydroxyphenyl 4-hydroxybenzoate (HPHB), 4-(Methoxymethyl) phenol (4MOP), *p*-hydroxybenzoic acid (PHBA), and 1,4-hydroquinone (HQ) were purchased from Shanghai Macklin Biochemical Technology Co., Ltd (Shanghai, China). BPF degradation experiments were performed in a mineral salts medium (MSM, pH 7.0). The bacterial strains used in this study were cultured in lysogeny broth (LB, pH 7.0) [21].

### Strains, plasmids, and cultural conditions

Details of the bacterial strains and plasmids utilized in this study are provided in Supplementary Table S1. *Microbacterium* sp. F2 was cultured in LB broth or MSM at 30℃ and 180 rpm, unless otherwise specified. *Escherichia coli* strains were cultured in LB broth at 37℃ and 200 rpm.

### BPF degradation assay

Strain F2 was initially cultured in LB medium for 24 h, followed by centrifugation at 4°C (5,000 × *g*, 5 min). The harvested cells were washed three times with sterile MSM. The cells, adjusted to an OD_600_ of 1.0, were then inoculated into MSM supplemented with 0.10 mM BPF and incubated at 30°C. Aliquots (1 mL) were taken every 2 h and analyzed via high-performance liquid chromatography (HPLC) to quantify BPF concentrations. A control experiment was conducted using MSM without strain F2 under identical conditions. All treatments were performed in triplicate.

To prepare BPF-induced cells, strain F2 cells cultured in LB medium were harvested, washed three times with sterile MSM, and inoculated into MSM containing 0.10 mM BPF. When half of the BPF was degraded, the BPF-induced cells were collected. For BPF-induced cell extracts, the cells were washed twice with 20 mM Tris-HCl buffer (pH 8.0), resuspended in 15 mL of the same buffer, and lysed using an ultrasonic processor (UH-650B, Auto Science) in an ice bath. The lysates were then centrifuged at 12,000 × g for 20 min at 4 °C. For the BPF degradation experiments, a 3 mL reaction mixture containing 0.10 mM BPF and 1 mL of cell lysate was prepared in a 20 mM Tris-HCl buffer (pH 8.0) at 30°C. The reaction was terminated immediately by boiling for 5 min after completion. Samples were collected every 10 min and analyzed by HPLC to measure the concentration of BPF.

### Sequencing and analysis

Genomic DNA from strain F2 was extracted using a commercial kit (TIANGEN BIOTECH Co., LTD, Beijing, China). The genome of strain F2 (accession number: JBNVBV000000000) was sequenced by Majorbio Biomedical Technology Co., Ltd (Shanghai, China). Gene prediction and annotation were carried out using Glimmer 3.02 and RAST (Rapid Annotation Subsystem Technology) database (https://rast.nmpdr.org/) [22].

### Purification of BpfA from the cell extract of strain F2

The purification was conducted at 0-4°C using a 20 mM Tris-HCl buffer (pH 8.0). The detailed protocol followed the methods previously reported [45]. The protein band visualized on SDS-PAGE (Fig. S3, lane 6) was excised and subjected to peptide mass spectrometry analysis. The resulting peptide fragments were matched against the annotated genes from strain F2 to pinpoint sequences with high homology. Subsequently, the resulting protein sequences underwent a BLASTP search against the NCBI non-redundant protein (NR) database. Based on these analyses, the gene encoding FAD-binding oxidoreductase, *bpfA*, was identified as a potential candidate for catalyzing BPF and was selected for further protein expression and *in vitro* BPF degradation assays.

### Bioinformatic Analysis

A BLASTP search was conducted using the BpfA sequence (PV658135) from *Microbacterium* sp. F2 as the query sequence, targeting all genomes within the NCBI database to uncover BpfA homologues. The sequence similarity network (SSN) was created utilizing the Enzyme Function Initiative Enzyme Similarity Tool (EFI-EST) [23]. An all-by-all BLAST was performed on the sequences, and the SSN was subsequently built based on an alignment score cutoff of 90. Visualization of the SSN was accomplished with Cytoscape [24]. For the phylogenetic analysis of BpfA, homologous sequences were retrieved from the NCBI NR database. These sequences were aligned using the ClustalX [25], and the resultant alignment was transferred to MEGA (version 11.0) [26] for further processing. The phylogenetic tree was subsequently generated employing the neighbor-joining algorithm.

Previously, species-level genome bins (SGBs) were constructed from 3,304 soil metagenomes, ensuring medium to high quality standards (with completeness exceeding 50% and contamination below 10%). Further details are available at the Global Soil MAGs Project website [27]. These SGBs were classified using GTDB-Tk (version 1.6.0), and the BpfA protein was identified through Prodigal, with stringent criteria (identity greater than 35%, e-value less than 1 × 10^−5^, and subject coverage over 95%) [28]. To explore the global distribution of BpfA, amino acid sequences from the SGBs were obtained and analyzed via Diamond (v2.1.6). By integrating classification data from SGBs and GTDB, a comprehensive analysis of BpfA species distribution was accomplished [29]. Additionally, the classification of target genomes as either isolated or uncultured (including metagenomic assembly genomes and single amplified genomes) was determined by searching both GTDB and NCBI RefSeq [30].

### Gene expression and purification of the recombinant enzyme

The primer pair *bpfA*_F2_-F and *bpfA*_F2_-R (Supporting Information Table S2) was used to amplify the *bpfA* gene from strain F2. The amplified *bpfA* gene was cloned into the NdeI/XhoI-digested pET29a(+) plasmid via homologous recombination, resulting in the recombinant plasmid pET-*bpfA*. This plasmid was then introduced into *Escherichia coli* BL21(DE3) to create the expression strain *E. coli* BL21(pET-*bpfA*). The strain *E. coli* BL21(pET-*bpfA*) was inoculated into LB containing 50 mg/L kanamycin and cultured until the cell density reached an OD_600_ of 0.5-0.6, at which point 0.2 mM IPTG was added to induce protein expression for 12 h at 16°C. Cells were centrifuged and lysed using an ultrasonic processor. The supernatant was obtained by centrifugation at 4°C and then passed through a Ni-NTA column (Sangon, Shanghai, China). A series concentrations of imidazole were used to elute the column and purify the recombinant protein. The purified protein was then dialyzed overnight in Tris-HCl buffer (pH 8.0) and analyzed by SDS-PAGE. A BCA kit from Vazyme Biotech Co., Ltd (Nanjing, China) was used to determine the protein concentration.

The genes *eugO* from *Rhodococcus jostii* RHA1 (Q0SBK1), *vaO* from *Penicillium simplicissimum* CBS 170.90 (P56216), and FAD-binding oxidoreductase gene *fbO* from *Sphingobium* sp. W15 (PV658134) were synthesized from Nanjing Qingke Biotechnology Co., Ltd (Nanjing, China). The protein expression and purification methods for EUGO, VAO, and FAD-binding oxidoreductase were similar to those described above.

### Enzyme activity assays

Enzymatic assays for BpfA were performed at 30°C for 2 h in a 1 mL reaction mixture containing 20 mM Tris-HCl buffer (pH 8.0), 0.10 mM BPF, and 3.0 μM BpfA. The enzymatic reactions were terminated by boiling for 5min. Enzyme activity was quantified as the amount of enzyme needed to convert 1 μmol of BPF per minute, defined as one unit.

To determine the optimal reaction temperature, BpfA’s activity was tested at various temperatures, spanning from 20°C to 50°C in 5°C increments. The thermal stability of BpfA was evaluated by subjecting the enzyme to different temperatures for 30 min, after which the residual activity was assessed under the specified assay conditions. The non-heated enzyme was used as the control (100%). The optimal reaction pH was determined using a variety of buffers: 20 mM Citrate-Phosphate buffer (pH 4.0 to 6.0), 20 mM Tris-HCl buffer (pH 6.0 to 8.0), and 20 mM glycine-NaOH buffer (pH 8.0 to 10.0). The pH stability of BpfA was examined by incubating the enzyme at 4°C for 30 min in buffers with diverse pH, with residual activities subsequently measured. Samples were collected prior to complete substrate consumption. A reaction mixture devoid of the enzyme served as a negative control. The activity of the reference enzyme was designated as 100%, and the relative activities of each reaction were determined as a percentage relative to the reference.

To evaluate the substrate specificity of BpfA, the enzyme was tested with various substrates (BPE, VA, 4PG, and 4MOP) at a concentration of 0.10 mM each, under the reaction conditions described above. The activity assays for EUGO, VAO, and FBO were similarly performed following the same procedures.

### Site-directed mutagenesis

Mutations at specific sites within the *bpfA* gene were engineered via overlap extension PCR. The primer pairs Y93A-F/R, Y474A-F/R, R475A-F/R, D152A-F/R, and H393A-F/R were used to introduce the mutation sites (Table S2). The site-directed mutagenesis amplification was carried out following the protocol provided with the site-directed mutagenesis kit (Vazyme Biotech Co., Ltd., China). The resulting PCR products were purified by gel extraction and ligated into pET-29a(+) vector, following the previously described method. The activity assessment of the mutant proteins was carried out using the previously detailed procedures.

### Analytical methods

Enzyme assay samples were boiled for 5min and subsequently centrifuged at 12,000 × *g* for 5 min. The resulting supernatant was passed through a 0.2-mm-pore-size membrane prior to further analysis of the products. The detection and quantification of BPF and its metabolites in enzyme assay samples were performed using HPLC [21]. The identification of BpfA and its metabolites in enzyme assay samples was achieved using mass spectrometry on an AB SCIEX Triple TOF 5600 [21].

## Results and discussion

### Cloning and sequence analysis of *bpfA* gene

*Microbacterium* sp. F2 has been reported to use BPF as the sole carbon source for growth, with BPF degradation being induced by BPF itself [21]. Furthermore, cell extract from BPF-induced strain F2 completely degraded 0.10 mM BPF, regardless of whether NADPH, NADH, and FAD were introduced (Fig. S2). Therefore, the gene encoding the initial step of BPF degradation can be obtained from the cell extract of strain F2 via protein purification techniques. A four-stage purification protocol was utilized to refine the BPF-degrading enzyme from the cell extract of BPF-induced strain F2. Initially, the specific activity in the cell extract from strain F2 was determined to be 0.37 U/mg protein. Following the four purification steps, the specific activity of the final fraction rose to 14.74 U/mg, with a 24.15-fold increase in protein concentration and a recovery rate of 4.76% (Table S3). Subsequent SDS-PAGE analysis of the final fraction showed a single protein band (Fig. S3), which was excised and subjected to peptide mass spectrometry analysis. The analysis yielded sequences of fifteen peptide fragments, which were then matched against all annotated genes within strain F2 (JBNVBV000000000) (Table S4). Subsequently, all candidate genes predicted to degrade BPF were annotated and experimentally validated. A FAD-binding oxidoreductase-encoding gene, *orf1733* (ACQVDU_06965), demonstrated BPF-degrading activity and was designated as *bpfA*, which was selected for further studies. The *bpfA* gene (1,599 bp) encodes a 532-amino acid oxidase that shares 30%–55% identity with FAD-binding oxidoreductases from various genera (Non-redundant Protein Sequences Database). BpfA is classified within the flavoprotein oxidase superfamily [31,32]. A sequence similarity network (SSN) analysis indicates that BpfA is a member of the vanillyl alcohol oxidase (VAO) flavoprotein family (PF01565) within this superfamily (Fig. 1A). BpfA shows 33% to 42% sequence identity with three biochemically characterized oxidoreductases from *Rhodococcus jostii* RHA1 (vanillyl-alcohol oxidase EUGO [Q0SBK1]; 41.14% identity) [33], *Penicillium simplicissimum* CBS 170.90 (vanillyl-alcohol oxidase VAO [P56216]; 37.50% identity) [34], and *Pseudomonas putida* NCIMB 9869 (*p*-cresol methylhydroxylase PCMH [P09788.3]; 33.20% identity) [35]. These three enzymes are members of the VAO flavoprotein family, which is characterized by a conserved FAD-binding domain [36] and a Tyr-Tyr-Arg triad essential for substrate binding [37]. Additionally, the VAO flavoprotein family (PF01565) is further divided into 11 subfamilies [37]. BpfA clusters with the reported members of the 4-phenol oxidizing (4PO) subfamily, including EUGO (Q0SBK1), PCMH (P09788.3), and VAO (P56216), suggesting that these proteins may have a closer phylogenetic relationship and similar functions (Fig. 1B). BpfA and other members of the 4PO subgroup, including EUGO, PCMH, and VAO, share five conserved motifs (Fig. S4) [38]. The residues in the 4PO-motif-1 form a loop that closely interacts with the FAD cofactor. Notably, one of the tyrosine residues (Tyr93), which is essential for substrate deprotonation, is situated within this loop. The 4PO-motif-5 also forms a loop that includes the other two residues, Tyr474 and Arg475, involved in substrate deprotonation. Furthermore, the amino acids Asp152 and His393 are involved in forming a covalent bond with FAD, which is critical for the enzyme’s redox properties. This covalent binding significantly enhances the redox potential of the enzyme [39], thereby accelerating the rate of FAD reduction by the substrate and, consequently, enhancing the overall reaction rate [39].

**Fig. 1.**
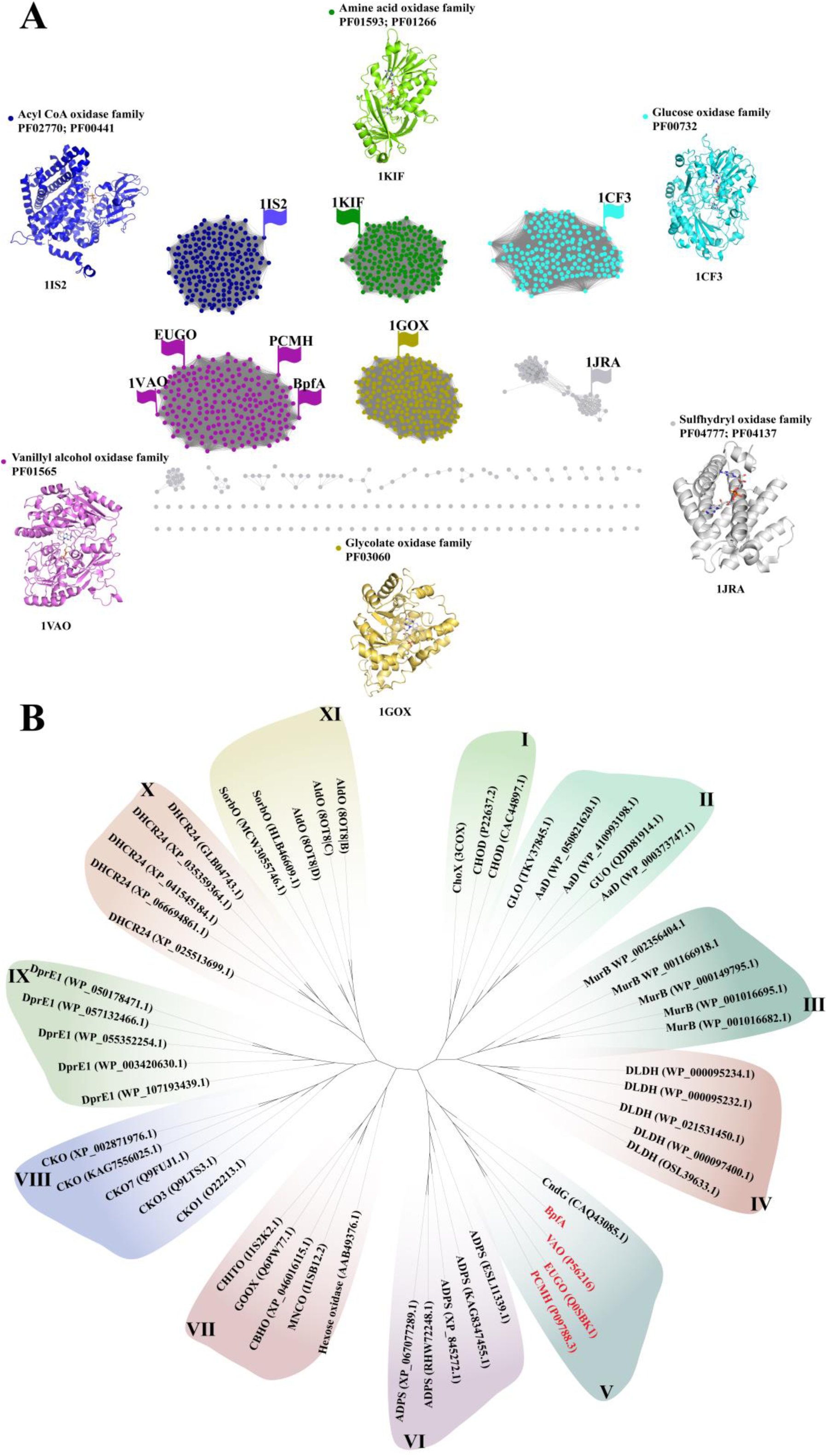
(A) Sequence similarity network (SSN) for representative flavoprotein oxidase superfamily in the NCBI Non-redundant Protein Sequences Database. Each item is arranged in the following order: protein name and accession number. The around are shown the structures of representative members of the different flavin-dependent oxidases with the FAD or FMN cofactor highlighted in gold. (B) Phylogenetic analysis of BpfA with the sequences of other members of the VAO flavoprotein family. Ⅰ: cholesterol oxidases subfamily; Ⅱ: aldonolactone oxidoreductases subfamily; Ⅲ: MurBs subfamily; Ⅳ: α-Hydroxy acid dehydrogenases subfamily; Ⅴ: 4-phenol oxidizing subfamily; Ⅵ: alkyl-dihydroxyacetone phosphate synthases subfamily; Ⅶ: BBE-like subfamily; Ⅷ: cytokinin dehydrogenases subfamily; Ⅸ: DprE1s subfamily; Ⅹ: sterol reductases subfamily; XI: alditol oxidases subfamily.

### Functional identification of BpfA

The purified BpfA protein displayed a pale yellow and showed a single band on SDS-PAGE, with an observed molecular mass of 59.9 kDa, which closely aligns with the theoretical molecular mass of 58.8 kDa (Fig. 2A). Purified BpfA was able to completely degrade 0.10 mM of BPF *in vitro* within 40 min. During the degradation of BpfA, four metabolites were identified (namely compound I, II, III, and IV) (Fig. 2B). The peak height of compound I (5.000 min) continuously decreased and completely disappeared at 40 min. After 20 min, compound II (3.998 min), compound III (2.708 min), and compound IV (3.703 min) appeared simultaneously with the degradation of compound I. However, compound II only appeared briefly before rapidly disappearing. Subsequently, compound III was gradually converted to compound IV and completely disappeared after 60 min. Ultimately, the peak height of compound IV continuously increased and remained unchanged. These compounds were identified by HPLC-MS as BPF (compound I, Fig. 2C), 4-[(4-hydroxyphenyl)methylene]cyclohexa-2,5-dien-1-one (HMC) (compound II, Fig. 2D), BHPM (compound III, Fig. 2E), and DHBP (compound IV, Fig. 2F). Additionally, we detected the production of 0.189 mM hydrogen peroxide (H_2_O_2_) during the degradation of 0.10 mM BPF by BpfA using the H_2_O_2_ colorimetric method [40], with a ratio of approximately 1:2 (Fig. S5).

**Fig. 2.**
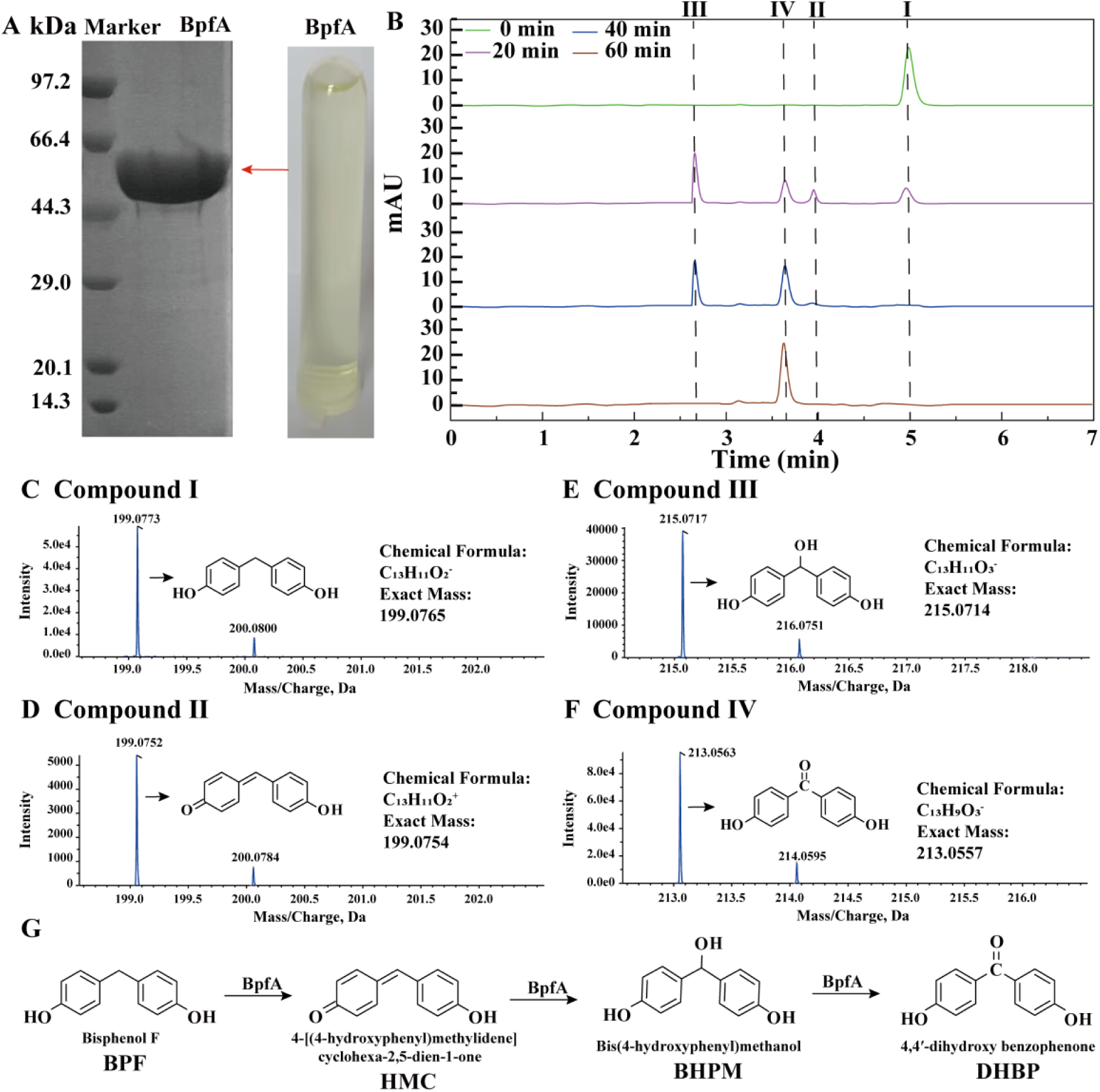
HPLC-MS analysis of BPF products treated with BpfA. (A) SDS-PAGE analysis of the purified BpfA. (B) HPLC analysis of metabolites that appeared during the conversion of BPF by purified BpfA. (C) HPLC-MS analysis of compound I (*m/z* 199.0773 [M-H]^−^), which was identified as BPF. (D) HPLC-MS analysis of compound II (*m/z* 199.0752 [M+H]^+^), which was identified as HMC. (E) HPLC-MS analysis of compound Ⅲ (*m/z* 215.0717 [M-H]^−^), which was identified as BHPM. (F) HPLC-MS analysis of compound Ⅳ (*m/z* 213.0563 [M-H]^−^), which was identified as DHBP. (G) The proposed metabolic pathway of BPF degradation by BpfA. BPF: bisphenol F; HMC: 4-[(4-hydroxyphenyl)methylidene]cyclohexa-2,5-dien-1-one; BHPM: bis (4-hydroxyphenyl) methanol; DHBP: 4,4’-dihydroxybenzophenone.

The identified metabolites led to the proposal of a BPF degradation pathway mediated by BpfA (Fig. 2G). Initially, the phenolic -OH group of BPF is deprotonated by two conserved tyrosine residues of BpfA (Y93 and Y474). The electron density is then conducted through the aromatic ring, extruding a hydride ion (H^−^) from the α-position, which is accepted by the isoalloxazine ring of the oxidized flavin cofactor. The resulting quinone methide intermediate (HMC) then acts as an electrophile, undergoing nucleophilic addition with a water molecule catalyzed in the BpfÁs active site, leading to the formation of the hydroxylated product (BHPM). Finally, BHPM undergoes further dehydrogenation to generate DHBP. Meanwhile, FAD is reoxidized by molecular oxygen, which is converted to H_2_O_2_. The initial deprotonation of the phenolic hydroxyl group of the substrate is a crucial step in this mechanism, as it facilitates the hydride transfer reaction by promoting the formation of the *para*-quinone methide intermediate. The same effect has been observed in VAO and EUGO [41,42].

In this study, BpfA catalyze the sequential transformation of BPF into the quinone methide intermediate HMC, BHPM, and DHBP (Fig. 2), which differs from the BPF degradation pathway proposed in strain F2, in which BPF was metabolized into BHPM and DHBP without the detection of the intermediate HMC [21]. According to the previously reported VAO catalytic mechanism, the key product intermediate, quinone methide, is transiently formed and allows for the rapid occurrence of several different oxidation reactions [41,42]. We speculate that the quinone methide intermediate HMC may be rapidly converted to BHPM in strain F2, and thus cannot be captured by HPLC. In this study, purified BpfA is used to catalyze BPF *in vitro* and detected HMC through multiple consecutive samplings. In short, The proposed BPF degradation pathway in strain F2 was described at the cellular level but lacked genetic or enzymatic evidence. However, this study elucidates the BPF degradation process at the enzymatic and genetic levels and proposes a modified BPF metabolic pathway compared to that by Wang et al. [21].

### BpfA and 4PO subfamily proteins

To investigate the potential distribution of BpfA across different strains, we searched the NCBI NR database for homologs with a coverage of no less than 95% and at least 30% amino acid sequence identity. Consequently, 49 gene pairs belonging to the 4PO subfamily were identified from three bacterial phyla, specifically from Pseudomonadota (12), Actinomycetota (27), and Ascomycota (10) (Fig. 3A). The BpfA homologs originate from soil, sediments, clinical specimens, wastewater, faeces, pulping wastewater, oil-well water-associated microorganisms, indicating their ubiquity in the environment. Phylogenetic tree analysis shows that BpfA, EUGO, VAO, and PCMH are located on different branches, suggesting that they may not have evolved directly from a single common ancestor but rather may have undergone different branching events during the course of evolution. In order to determine if these BpfA homologous genes were functional, we synthesized three genes from Ascomycota, Actinomycetota and Pseudomonadota: EUGO (Q0SBK1; *Rhodococcus jostii* RHA1; coverage 98%; identity 41.14%), VAO (P56216; *Penicillium simplicissimum* CBS 170.90; coverage 96%; identity 37.50%), and FAD-binding oxidoreductase FBO (PV658134; *Sphingobium* sp. W15; coverage 97%; identity 39.46%), for the purpose of expressing recombinant proteins in *E. coli*. Given that PCMH from *Pseudomonas putida* NCIMB 9869 (Pseudomonadota) is a two-component protein with an *α*2*β*2 structure, comprising an *α*-subunit (PchF) and a *β*-subunit (PchC) to function [35], and considering that the reported EUGO and VAO are both single-component proteins, this study did not consider synthesizing PCMH for subsequent functional validation.

**Fig. 3.**
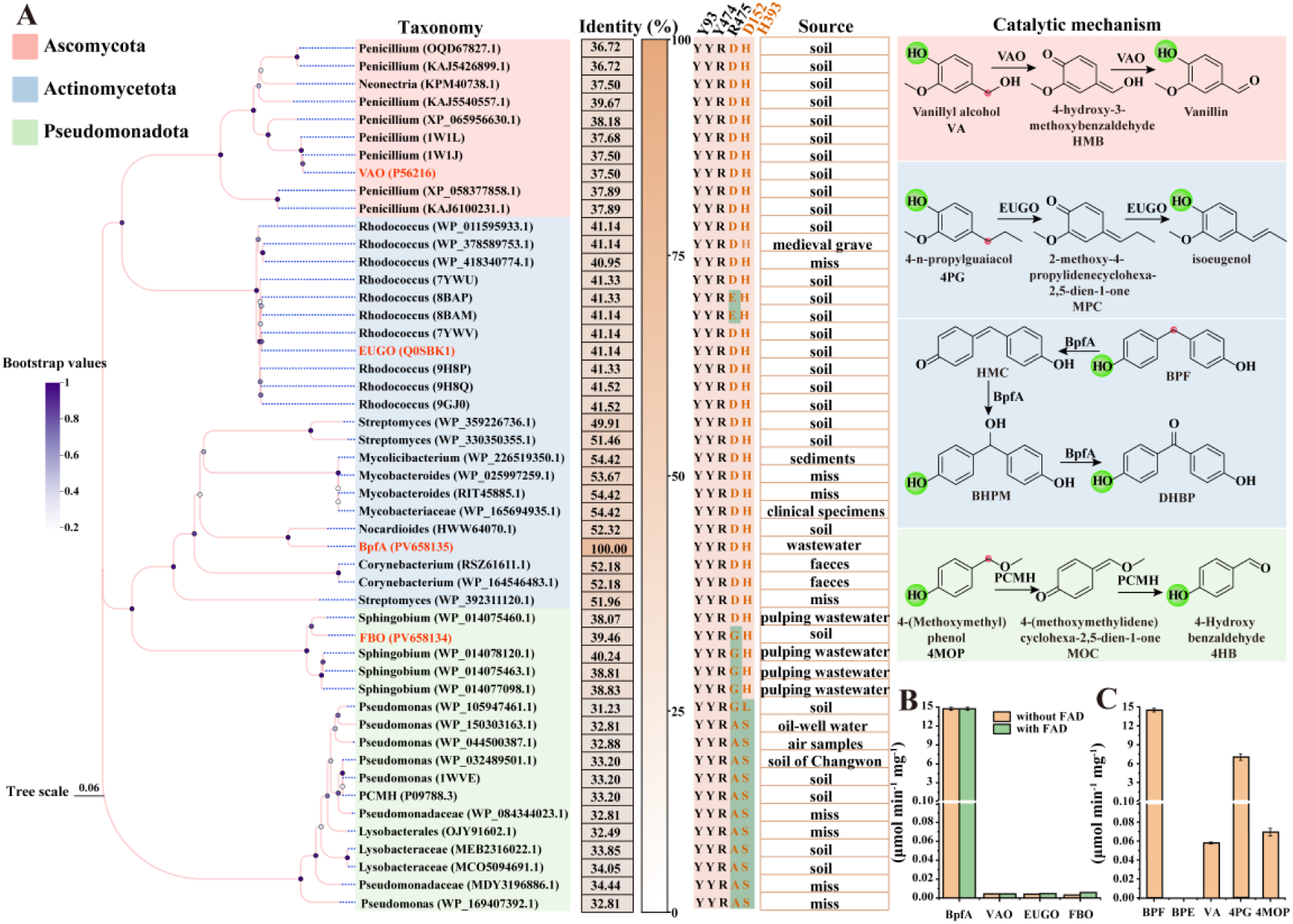
Bioinformatic and functional analyzes of the 4-phenol oxidizing (4PO) subfamily proteins. (A) Phylogenetic tree of BpfA. The identity of BpfA homologous sequences is indicated on the right side of the phylogenetic tree. The three conserved key amino acid residues (black) and two conserved FAD-binding sites (orange) for each sequence are shown on the right side, and their positions in BpfA are indicated on the top of the columns. The residues with a green background are mutated residues, differing from the conserved residues marked with a pink background. The sources of strains containing BpfA homologous sequences are listed on the right side of the conserved amino acid sequences. On the far right side of the phylogenetic tree detail the catalytic mechanisms of VAO, BpfA, EUGO, and PCMH for their respective substrates. Green spheres represent invariant residues; red spheres indicate the catalytic sites of the reactions. (B) Assessment of the BPF activity of BpfA, VAO, EUGO, and FBO. The data represent the mean values of two replicates using independent enzyme preparation. (C) Assessment of the BPF, BPE, VA, 4PG, and 4MOP enzyme activity of BpfA. The data represent the mean values of two replicates using independent enzyme preparation.

In the absence of FAD, VAO, EUGO, and FBO exhibited weak activity against BPF, with the specific activity of BpfA being 2452, 2907, and 3672 times higher than that of VAO, EUGO, and FBO, respectively (Fig. 3B). In the presence of FAD, the specific activity of BpfA was 2381, 2809, and 2104 times higher than that of VAO, EUGO, and FBO, respectively (Fig. 3B). There was no significant difference in the catalytic activity of VAO and EUGO against BPF, regardless of the presence or absence of FAD. Compared with VAO and EUGO, the addition of FAD significantly enhanced the catalytic activity of FBO against BPF. Further analysis revealed that FBO (D to G mutation) each had only one amino acid residue involved in FAD binding altered (Fig. 3A). Therefore, the addition of FAD was required for FBO to exhibit higher catalytic activity. The above results indicate that members of the 4PO subfamily proteins exhibit catalytic activity against BPF.

### Biochemical characterization of BpfA

The specific BpfA activity was 14.71 U mg^−1^ protein for BPF (Fig. 3B). The highest BPF activity was observed at a temperature of 30°C and a pH of 8.0 (Fig. S6). When the enzyme was exposed to 40°C for 30 min, it maintained over 80% of its initial activity. In contrast, incubation at 45°C for the same duration resulted in significant instability, with the enzyme retaining only 30% of its initial activity (Fig. S6A). BpfA exhibits high activity at a pH of 8.0, with its activity dropping below 20% at pH less than 5.0. When stored for 30 min at pH ranging from 7.0 to 10.0, its activity remains at 70% (Fig. S6B).

BpfA exhibits catalytic activity against other *para*-substituted phenolic compounds, including vanillyl alcohol (VA), 4-n-propylguaiacol (4PG), and 4-(methoxymethyl)phenol (4MOP) (Fig. 3C). The specific activity of BpfA against BPF is 249, 2.09, and 210 times higher than that against VA, 4PG, and 4MOP, respectively. However, BpfA does not exhibit catalytic activity against bisphenol E (BPE), which has a structure similar to BPF (Fig. S7), indicating that the two active hydrogen atoms on the α-position carbon of *para*-substituted phenolic compounds are crucial for BpfA to function.

Moreover, the Tyr-Tyr-Arg triad and conserved FAD-binding domain (Y93, Y474, R475, D152, and H393) of BpfA was replaced by alanine, resulting in five variants (BpfA^Y93A^, BpfA^Y474A^, BpfA^R475A^, BpfA^D152A^, and BpfA^H393A^) (Fig. S8). Among the five mutants, only the purified BpfA^Y474A^ and BpfA^H393A^ exhibited the same pale yellow as the wild-type BpfA (Fig. S8A). BpfA^Y474A^ and BpfA^H393A^ retained 6.6% and 71.1% of their activity against BPF, respectively, while the other mutants completely lost their catalytic activity against BPF (Fig. S8B). This indirectly highlights the significance of the Tyr-Tyr-Arg triad for BpfÁs functionality and also indicates that Asp152 plays a more crucial role than His393 in forming a covalent bond with FAD. This also explains why FBO require the addition of FAD to exhibit stronger catalytic activity. The results confirm that BpfA belongs to the 4PO subfamily, featuring the Tyr-Tyr-Arg triad and a conserved FAD-binding domain.

### Global survey on the environmental distributions of *bpfA* gene

Considering that BpfA homologs are predominantly derived from soil microbes (Fig. 3), we conducted a mapping of BpfA reference sequences onto the species-level microbial genomes present in GTDB and SGBs, aiming to investigate the diversity of BpfA throughout the microbial tree of life. Among 153,143 microbial species across 12 phyla, BpfA homologs were identified in 1.08% (1652 genomes) of the species. The study revealed that 67.37% of the microbial species potentially harboring the *bpfA* gene remain uncultured (Fig. S9). BpfA was frequently found among Pseudomonadota (70.04%) and Actinomycetota (20.46%) in the bacterial phyla. However, BpfA was not observed in the archaea phyla (Fig. 4A). Notably, the genera *Paraburkholderia*, *Novosphingobium*, and *Sphingobium* emerged as major bacterial taxa containing BpfA, whereas only a few strains within the genus *Microbacterium* harbor this enzyme (Fig. 4B). Combining previously reported BPF-degrading strains, we can infer that bacteria carrying the BpfA and involved in BPF degradation are widely distributed across Pseudomonadota and Actinomycetota. These results expand our knowledge of the variety of bacteria carrying BpfA that participate in BPF biodegradation. Grasping the diversity and distribution of these bacteria is essential for elucidating their ecological importance and potential applications.

**Fig. 4.**
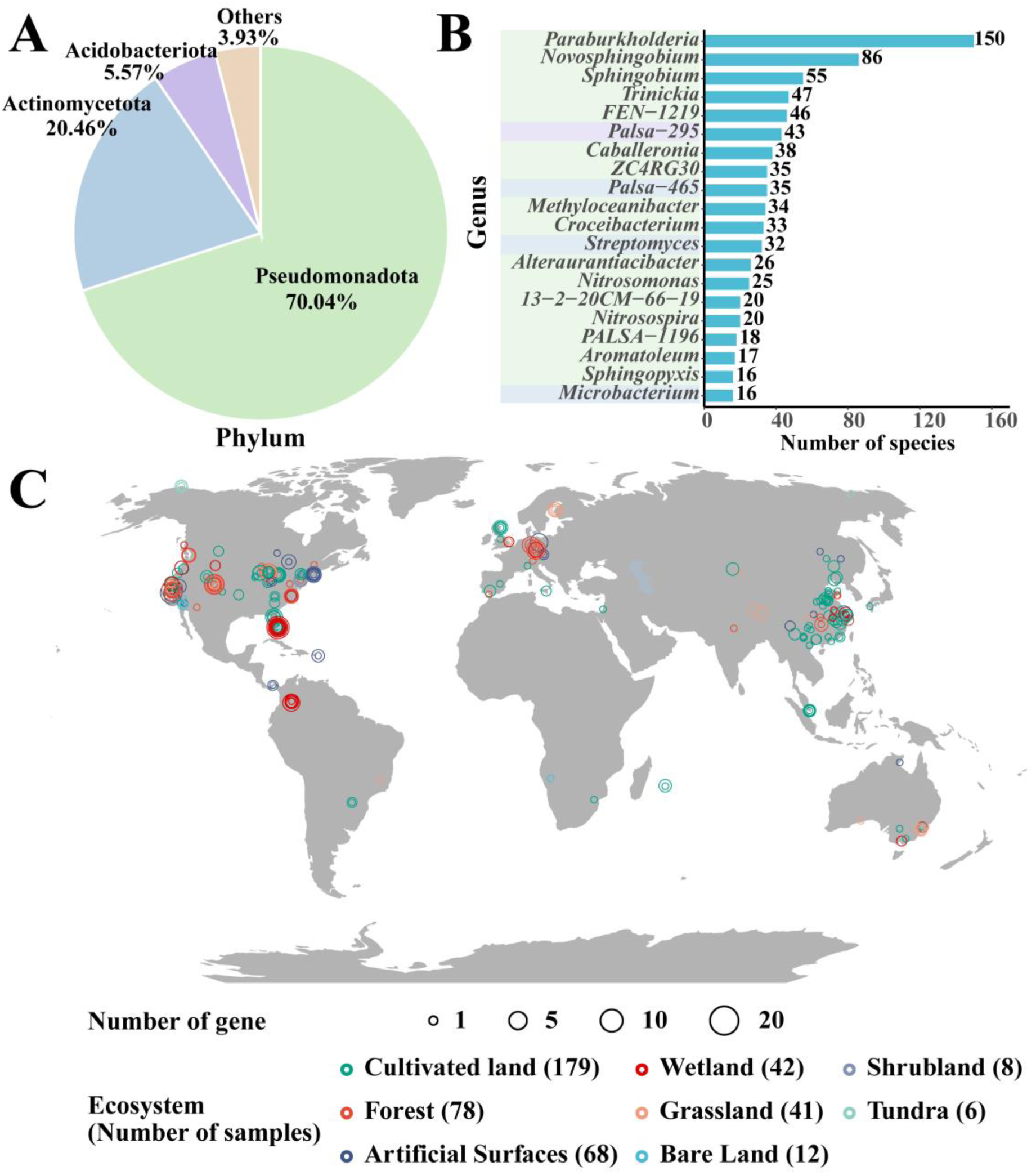
Microbial classification and distribution pattern of the BpfA in global environments. (A) The number of species that mapped the BpfA across phyla in the domain of bacteria and archaea. (B) The number of species that mapped the BpfA across genera in the domain of bacteria and archaea. (C) Geographic and abundance distribution of the BpfA-harboring bacteria.

The functional spectrum of BpfA was delineated by mapping its reference sequences with 3304 metagenomic samples from a wide range of global environments. The study found that BpfA exhibits distinct regional distribution patterns (Fig. 4C). Geographically, BpfA homologs is primarily distributed across Asia, North America, and Europe. These homologs are particularly abundant in environments such as cultivated land and forests, which produce large amounts of lignocellulosic waste (such as wheat straw, corn stover, and trees) annually. These wastes are degraded by soil microbes to produce 4-alkylphenols. The biodegradation of 4-alkylphenols in cultivated land and forests may be significantly influenced by bacteria carrying BpfA. In contrast, in environments with less vegetation and tree cover, such as bare land, shrubland, and tundra, the abundance of BpfA homologs is much lower.

Theoretically, the gene encoding BpfA-an enzyme capable of efficiently degrading BPF-may have existed in pre-industrial gene pools. In other words, the ancestral gene of *bpfA* could have been present in nature before humans synthesized BPF. This broad substrate specificity indicates that BpfA has evolved the capability to degrade a range of structurally similar compounds, likely stemming from its ancestral enzyme’s role in degrading naturally occurring 4-alkylphenols compounds. BpfA’s subsequent evolution to degrade the synthetic compound BPF supports the hypothesis that its progenitor existed pre-industrially to metabolize phenolic intermediates. However, it is also possible that there exists a relatively short evolutionary path from an unknown extant gene to the *bpfA* gene identified in current biodegradation pathways. It has been proposed that a novel sequence for an enzyme targeting a synthetic compound (e.g., BPF) could evolve via the activation of an unused alternative open reading frame within a pre-existing repetitive coding sequence [43]. However, it is important to note that the similarity between BpfA and members of enzyme superfamilies that catalyze other reactions typically does not provide specific insights into the adaptation process to xenobiotic compounds. The sequence similarity between BpfA and other proteins in phylogenetic families is generally less than 55% (Fig. 3), suggesting that the divergence of BpfA from these enzymes likely occurred much earlier than a century ago. Thus, its evolutionary process cannot be simply attributed to the introduction of industrial chemicals such as BPF into the environment. If BpfA and other key enzymes in metabolic pathways had undergone recent mutations, closely related sequences with only a few mutations differing from the current BpfA enzyme should be present in nature. However, to date, no such primitive BpfA enzyme has been found in the NCBI database. This suggests that BpfA may not have evolved through recent mutation events but rather through more complex evolutionary processes, such as gene mutation or recombination, to acquire its ability to degrade BPF.

Nevertheless, BpfA not only exhibits high degradative activity against BPF but also shows some activity towards 4-alkylphenols, which are products of lignocellulosic biomass reductive catalytic fractionation (RCF) [44], such as vanillyl alcohol (VA), 4-n-propylguaiacol (4PG), and 4-(methoxymethyl)phenol (4MOP). This broad substrate specificity of BpfA indicates that it has retained the metabolic function for natural phenolic compounds while evolving the capability to degrade BPF. It is precisely this enzymatic promiscuity of BpfA that makes it a potential biocatalyst for converting 4-alkylphenol compounds into valuable phenolic monomers.

## Conclusions

Given the widespread use of BPF, the residues of BPF in the environment cannot be overlooked. However, the metabolic degradation process of BPF by microorganisms is currently poorly understood. In this study, we cloned and identified a novel oxidase gene, *bpfA*, from *Microbacterium* sp. F2, which is responsible for catalyzing BPF into DHBP in a series of reactions. BpfA not only exhibits high degradation activity against BPF but also shows certain activity against 4-alkylphenol compounds such as VA, 4PG, and 4MOP. According to our survey of BpfA in the Global Soil MAGs Project metagenome database, BpfA homologs are widely distributed in the environment, especially in the metagenomes of cultivated land and forests, mainly belonging to Pseudomonadota and Actinomycetota. This study deepens our understanding of the microbial degradation mechanism of BPF, revealing the catalytic function of the novel oxidase gene *bpfA* and its distribution pattern in the environment.

## Author contributions

C.W., M.Z. and Q.H. conceived and developed the study. C.W., M.Z., Y.H. and Q.H. gathered the data and conducted the analyses. C.W., M.Z. and Q.H. led the writing of the manuscript. Q.L., J.H., K.P., Q.Z. and W.J. contributed critically to the analyses and writing. J.Q., X.Y. and Q.H. directed the study and critically revised the manuscript for important intellectual content. All authors edited the manuscript and approved the final version.

## Acknowledgements

This work was supported by the National Key R&D Program of China (2022YFA0912500), the National Natural Science Foundation of China (32170125 and 32400085), and the Postdoctoral Fellowship Program of CPSF (GZC20231127).

---

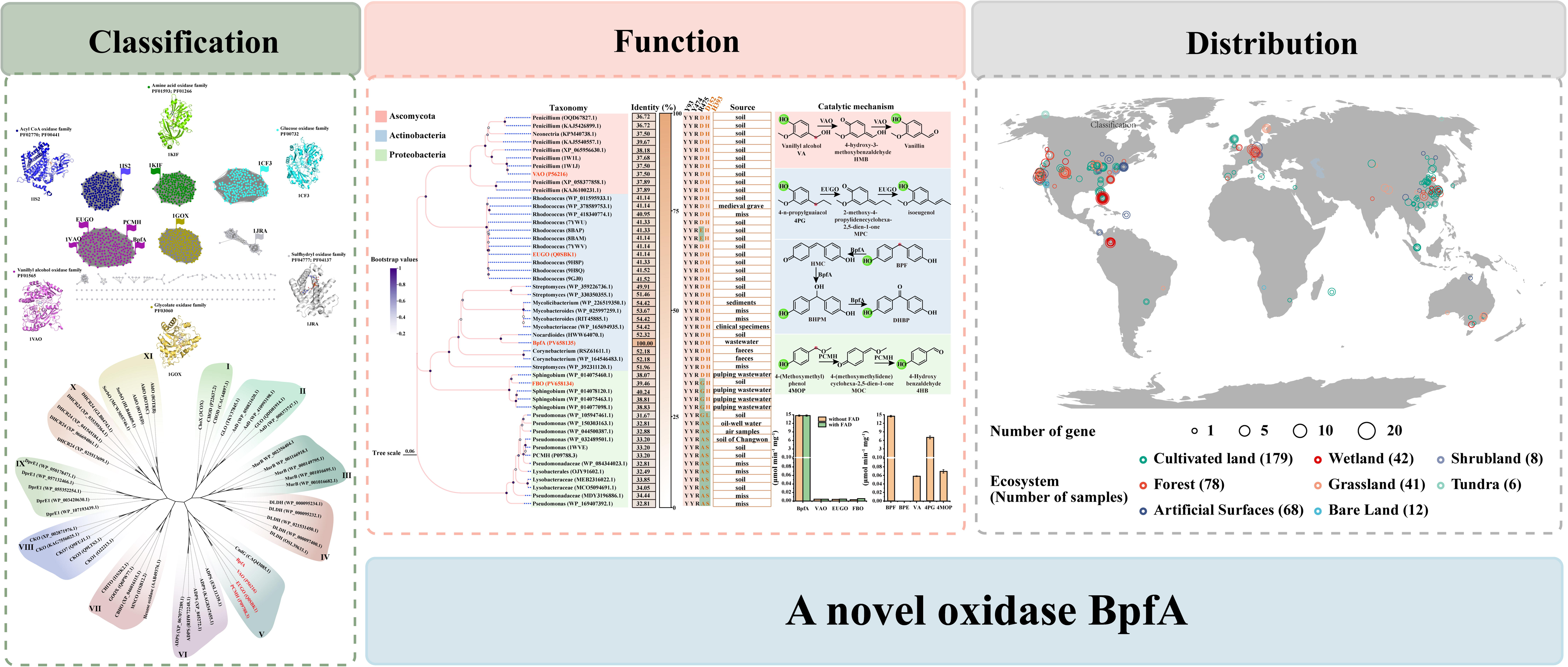

