## Supplemental Material for "Widespread distribution of the BpfA-carrying bacteria involved in the bisphenol F degradation in Pseudomonadota and Actinomycetota"

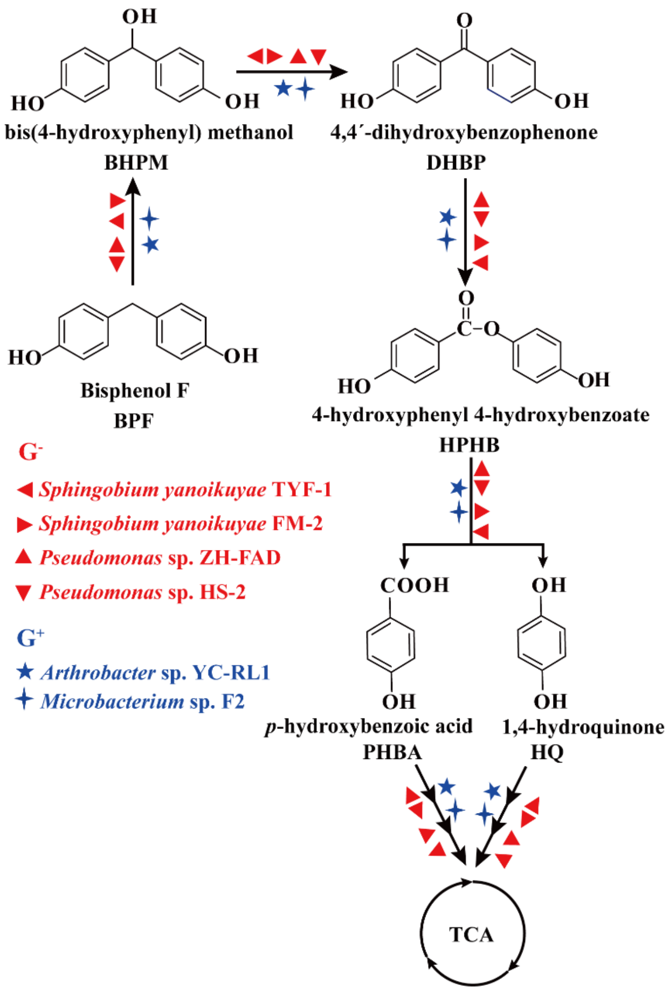


Fig S1. The conserved BPF degradation pathway in the reported strains. *Pseudomonas* sp. HS-2 [16], *Pseudomonas* sp. ZH-FAD [17], *Sphingobium yanoikuyae* FM-2 [18], *Sphingobium yanoikuyae* TYF-1 [19], *Arthrobacter* sp. YC-RL1 [20], and *Microbacterium* sp. F2 [24].

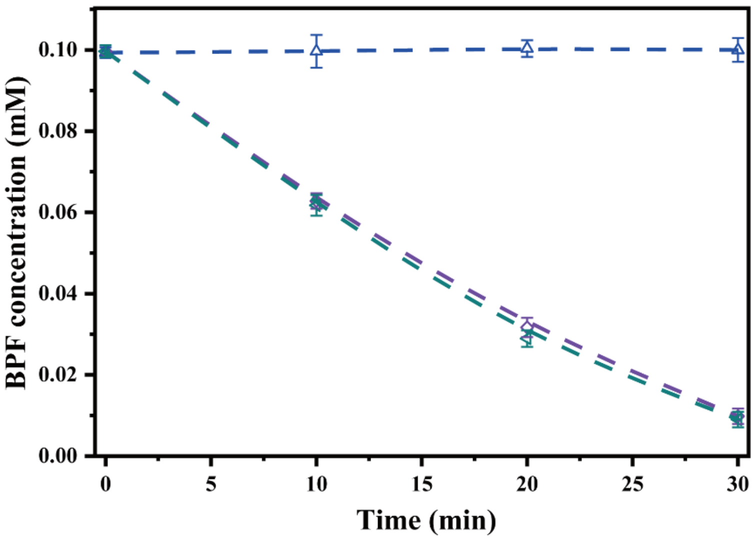


Fig S2. BPF degradation by the cell extracts of strains F2. △: Control; ◇: cell extract of strain F2; ◁: cell extract of strain F2 with FAD, NADPH, and NADH. Error bars represent the standard error of three replicates.

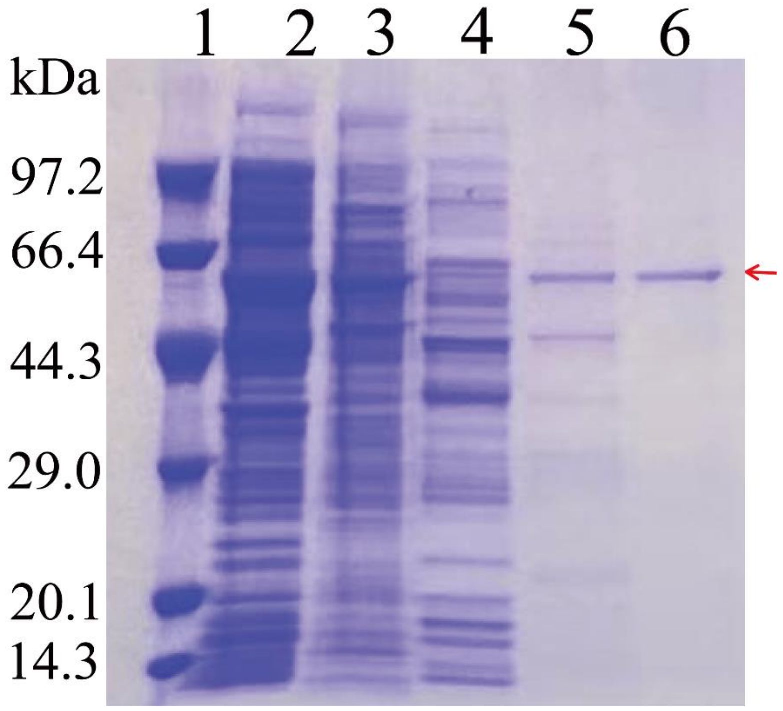


Fig S3. The purification of BpfA from strain F2 on SDS-PAGE. Lane 1: protein marker, Lane 2: cell extract, Lane 3: after ammonium sulfate precipitation, Lane 4: after by DEAE-Sepharose fast flow anion exchange chromatography, Lane 5: after Q-Sepharose fast flow anion exchange chromatography, Lane 6: after concentration by Microcon centrifugal filters and Sephadex-200 gel chromatography. The protein band (red arrow) were then excised and analyzed by matrix-assisted laser desorption ionization-time-of-flight (MALDI-TOF) mass spectrometry. Cell extract and ammonium sulfate precipitation fraction were diluted 10 and 4 times, respectively.

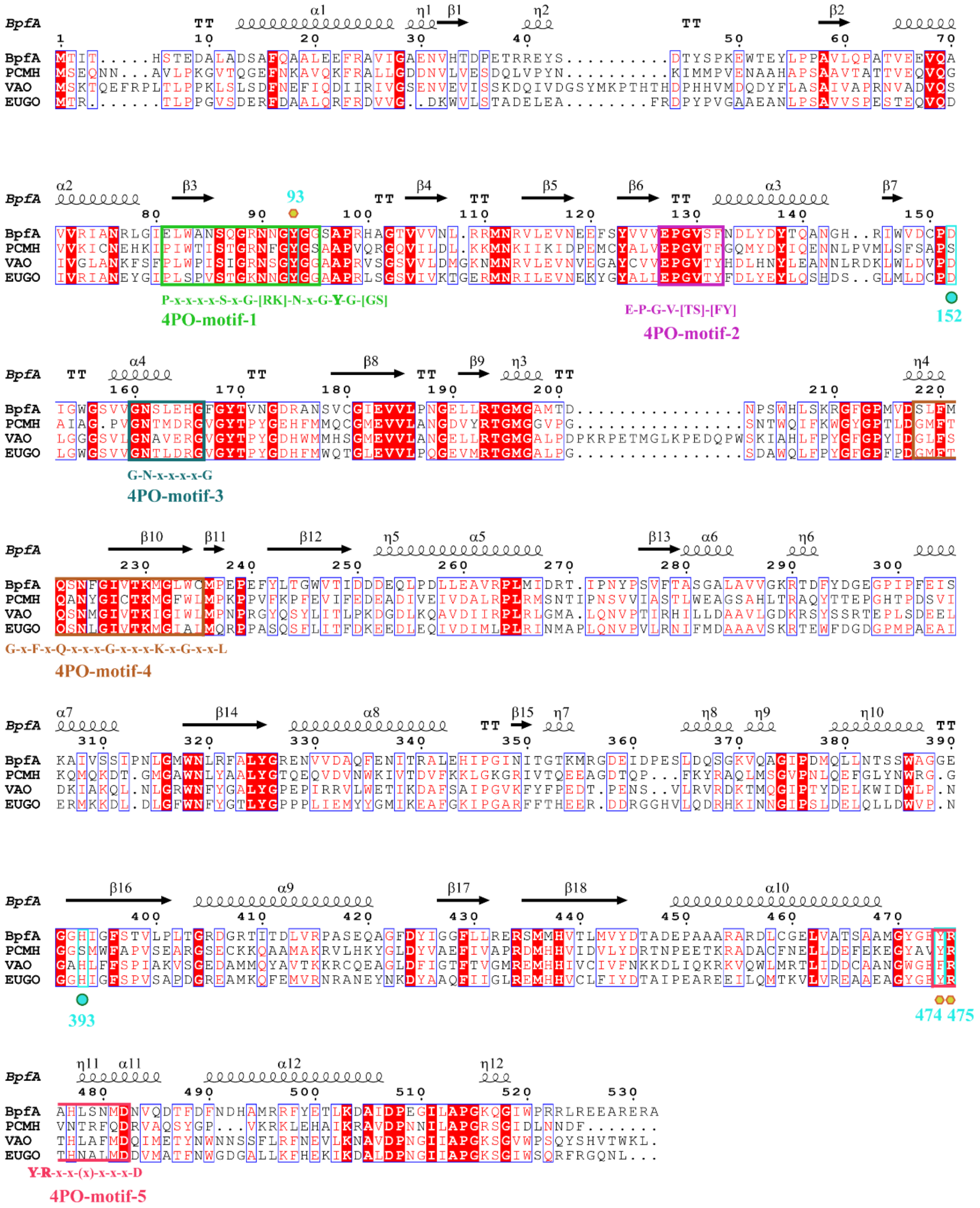


Fig S4. Sequence alignment of BpfA with the other members of the 4PO subgroup (EUGO, PCMH, and VAO). The five motifs of 4PO are labeled in different colors below the sequence. Bold letters indicate a critical amino acid residue. Pentagon represent the position of catalytic Tyr-Tyr-Arg (Tyr93, Tyr474, and Arg475) residues in BpfA. The FAD-binding site (Asp152 and His393) are denoted by circle.

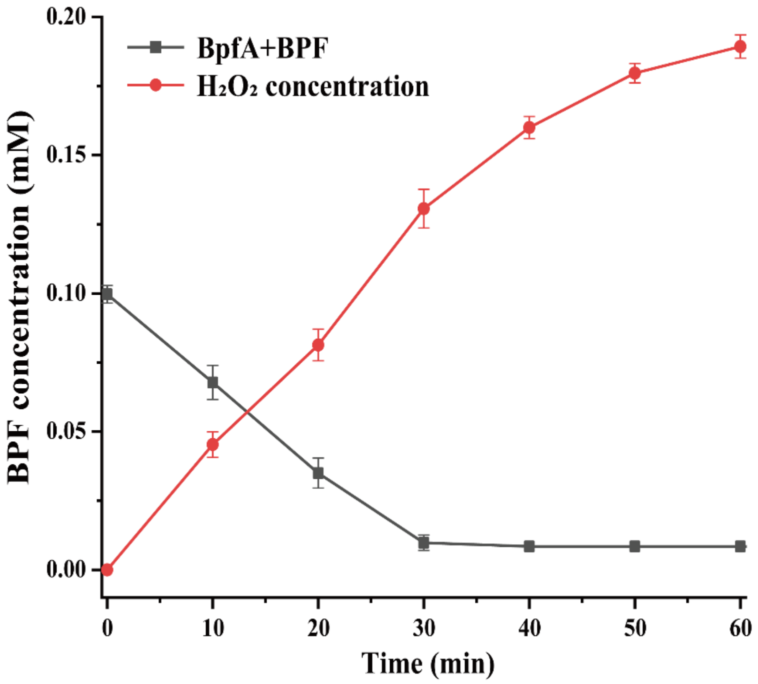


Fig S5. Hydrogen peroxide generated during the degradation of BPF. Black represents the degradation of BPF, and red represents the generation of $H_2O_2$.

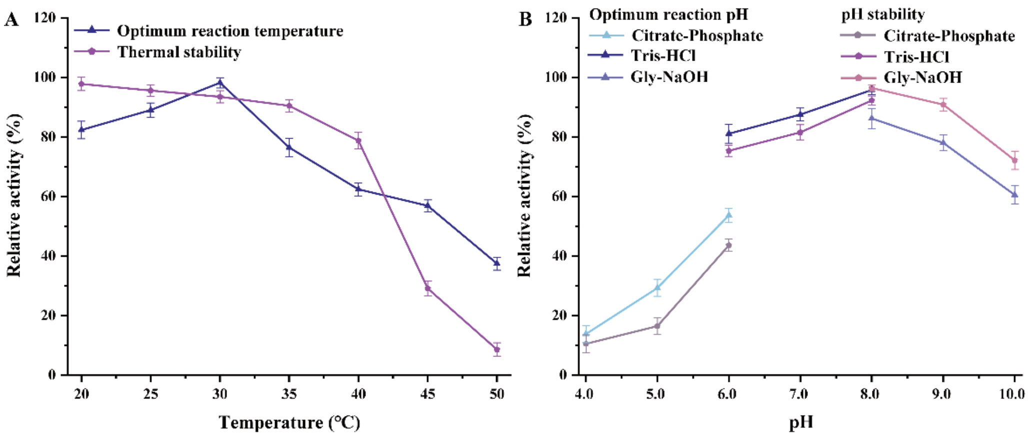


Fig S6. Effects of temperature and pH on the enzyme activity and stability of BpfA. The optimal temperature and thermal stability for the activities of BpfA (A); The optimal pH and pH stability for the activities of BpfA (B). The optimal temperature of BpfA (A) were determined using 20 mM Tris-HCl buffer (pH 7.0) at 20-50°C. The optimal pH of BpfA (B) was determined using 20 mM disodium hydrogen phosphate-citric acid buffer (pH 4.0 to 6.0), 20 mM Tris-HCl (pH 6.0 to 8.0) and 20 mM glycine-NaOH buffer (pH 8.0 to 10.0). The reaction without enzyme was used as a blank control. Error bars represent the standard error of three replicates.

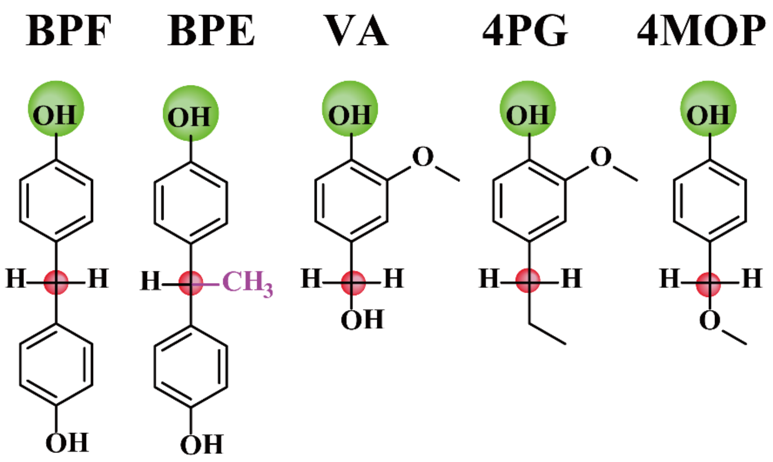


Fig S7. The chemical structure of 4-alkylphenols. Green spheres represent conserved chemical groups; red spheres indicate the catalytic sites of the reactions. bisphenol F (BPF); bisphenol E (BPE); vanillyl alcohol (VA); 4-*n*-propylguaiacol (4PG); 4-(Methoxymethyl) phenol (4MOP).

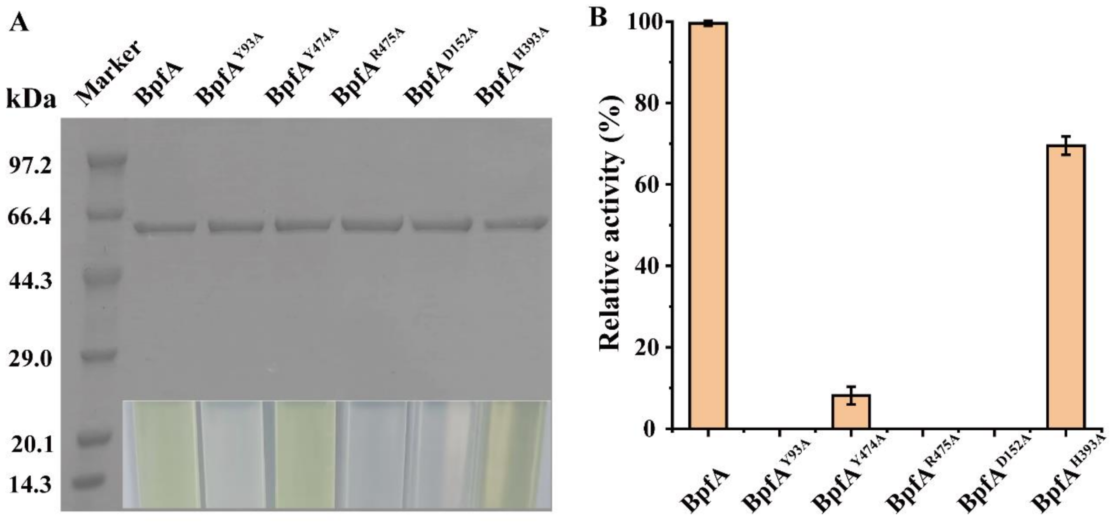


Fig S8. (A) SDS-PAGE analysis of the purified BpfA. Lane 1: protein marker; Lane 2: BpfA$_{F2}$; Lane 3: BpfA$_{F2}^{Y93A}$; Lane 4: BpfA$_{F2}^{Y474A}$; Lane 5: BpfA$_{F2}^{R475A}$; Lane 6: BpfA$_{F2}^{D152A}$; Lane 7: BpfA$_{F2}^{H393A}$. (B) Assessment of the BPF enzyme activity of BpfA$_{F2}$, BpfA$_{F2}^{Y93A}$, BpfA$_{F2}^{Y474A}$, BpfA$_{F2}^{R475A}$, BpfA$_{F2}^{D152A}$, and BpfA$_{F2}^{H393A}$. The data represent the mean values of two replicates using independent enzyme preparation.

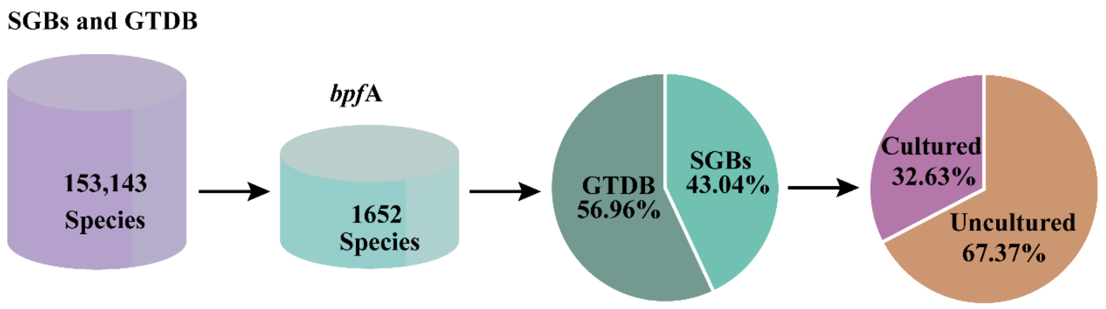


Fig S9. Species profiles with the BpfA from species-level genome bins (SGBs) and the Genome Taxonomy Database (GTDB).

Table S1 Strains and plasmids used in this study.

| Strains or plasmids | Description | Source |
|---|---|---|
| Strains | | |
| *Microbacterium* sp. F2 | Degrades bisphenol F; Gram-positive, Wild type | This study |
| *E. coli* strains | | |
| BL21(DE3) | F$^-$ *ompT hsdS*$_B$ ($r_B^-$ $m_B^-$) *dcm gal* λ(DE3) | Vazyme |
| Plasmids | | |
| pET-29a(+) | Km$^r$; Expression vector | Lab stock |
| pET-*bpfA*$_{F2}$ | Km$^r$; *NdeI-XhoI* fragment containing *bpfA*$_{F2}$ gene inserted into pET29a(+) | This study |
| pET-*bpfA*$_{F2}^{Y93A}$ | Km$^r$; *NdeI-XhoI* fragment containing *bpfA*$_{F2}^{Y93A}$ gene inserted into pET29a(+) | This study |
| pET-*bpfA*$_{F2}^{Y474A}$ | Km$^r$; *NdeI-XhoI* fragment containing *bpfA*$_{F2}^{Y474A}$ gene inserted into pET29a(+) | This study |
| pET-*bpfA*$_{F2}^{R475A}$ | Km$^r$; *NdeI-XhoI* fragment containing *bpfA*$_{F2}^{R475A}$ gene inserted into pET29a(+) | This study |
| pET-*bpfA*$_{F2}^{D152A}$ | Km$^r$; *NdeI-XhoI* fragment containing *bpfA*$_{F2}^{D152A}$ gene inserted into pET29a(+) | This study |
| pET-*bpfA*$_{F2}^{H393A}$ | Km$^r$; *NdeI-XhoI* fragment containing *bpfA*$_{F2}^{H393A}$ gene inserted into pET29a(+) | This study |
| pET-*fbO*$_{W15}$ | Km$^r$; *NdeI-XhoI* fragment containing *fbO*$_{W15}$ gene inserted into pET29a(+) | This study |
| pET-*vaO*$_{CBS\ 170.90}$ | Km$^r$; *NdeI-XhoI* fragment containing *vaO*$_{CBS\ 170.90}$ gene inserted into pET29a(+) | This study |
| pET-*eugO*$_{RHA1}$ | Km$^r$; *NdeI-XhoI* fragment containing *eugO*$_{RHA1}$ gene inserted into pET29a(+) | This study |

Table S2 Primers used in this study.

| Primers | Sequence (5'–3') [a] | Description |
| --- | --- | --- |
| *bpfA*$_{F2}$-F | TAAGAAGGAGATATACATATGATGACCATCACCCATTCGACCGAGGACGC | To construct plasmid pET-*bpfA*$_{F2}$ |
| *bpfA*$_{F2}$-R | GTGGTGGTGGTGGTGCTCGAGTGCCCGCTCCCGCGCCTC | |
| Y93A-F | GCAGGGCCGCAACAACGGCGCCGGCGGGTCCGCCCCGCG | To construct plasmid pET-*bpfA*$_{F2}^{Y93A}$ |
| Y93A-R | GCGCGGGGCGGACCCGCCGGCGCCGTTGTTGCGGCCCTGCGAGTTGGC | |
| Y474A-F | GGCGATGGGCTACGGGGAGGCTCGCGCGCACCTGTCGAA | To construct plasmid pET-*bpfA*$_{F2}^{Y474A}$ |
| Y474A-R | GTTCGACAGGTGCGCGCGAGCCTCCCCGTAGCCCATCGC | |
| R475A-F | GATGGGCTACGGGGAGTATGCCGCGCACCTGTCGAACAT | To construct plasmid pET-*bpfA*$_{F2}^{R475A}$ |
| R475A-R | CATGTTCGACAGGTGCGCGGCATACTCCCCGTAGCCCATCG | |
| D152A-F | CATCTGGGTCGACTGCCCCGCCATCGGGTGGGGGAGCGT | To construct plasmid pET-*bpfA*$_{F2}^{D152A}$ |

| | | |
|---|---|---|
| D152A-R | CGACGCTCCCCCACCCGATGGCGGGGCAGTCGACCCAGATG | |
| H393A-F | GGCGGGCGGGGAGGGCGGCGCCATCGGCTTCTCGACCGTGCTGCCGC | To construct plasmid pET-*bpfA*$_{F2}^{H393A}$ |
| H393A-R | CACGGTCGAGAAGCCGATGGCGCCGCCCTCCCCGCCCGC | |
| *nofbO*$_{W15}$-F | TAAGAAGGAGATATACATATGATGATGTCCGAACACTTGCCC | To construct plasmid pET-*fbO*$_{W15}$ |
| *fbO*$_{W15}$-R | GTGGTGGTGGTGGTGCTCGAGCAGATCGCCTCCCTTCCC | |
| *eugO*$_{RHA1}$-F | TCCGTCGACAAGCTTGCGGCCGCACTCGAGATGACAAGGACTCTACCCCCAG | To construct plasmid pET-*eugO*$_{RHA1}$ |
| *eugO*$_{RHA1}$-R | ATCTCAGTGGTGGTGGTGGTGGTGCTCGAGTTAGAGGTTCTGACCGCGGAAAC | |
| *vaO*$_{CBS\ 170.90}$-F | GTGGTGGTGGTGGTGCTCGAGCAGTTTCCAGCCCACTTG | To construct plasmid pET-*vaO*$_{CBS\ 170.90}$ |
| *vaO*$_{CBS\ 170.90}$-R | TAAGAAGGAGATATACATATGATGTCCAAAGCCTCCCCG | |

Table S3 Purification of BpfA from strain F2

| Step | Total protein (mg) | Total activity (U) | Specific activity (U/mg) | Recovery (%) | Fold |
|---|---|---|---|---|---|
| Cell extract | 159.03 | 58.84 | 0.37 | 100 | 1 |
| Ammonium sulfate precipitation | 20.89 | 31.13 | 1.49 | 52.91 | 2.51 |
| DEAE-Sepharose chromatography | 8.51 | 20.59 | 2.42 | 34.99 | 6.34 |
| Q-Sepharose chromatography | 1.83 | 13.89 | 7.59 | 23.61 | 13.74 |
| Sephadex-200 gel chromatography | 0.19 | 2.80 | 14.74 | 4.76 | 24.15 |

Table S4 The result of peptide mass spectrometry analysis of protein band

| ORF no. (locus_tag) | Location | Peptides[a] | PSMs[b] | AAs[c] | MW [kDa] | Score[d] | Homologous protein[e] | GenBank accession no. | Identity (%) |
|---|---|---|---|---|---|---|---|---|---|
| 1733 (ACQVDU_06965) | 1404956-1406554 | 23 | 83 | 532 | 58.80 | 188.39 | FAD-binding oxidoreductase [*Mycolicibacterium*] | WP_226519350 | 54.42% |
| 1280 (ACQVDU_15455) | 3130960-3132243 | 24 | 82 | 427 | 445.20 | 170.48 | NAD(P)/FAD-dependent oxidoreductase [*Microbacterium*] | WP_292728035 | 100.00% |
| 1054 (ACQVDU_14355) | 2941356-2942783 | 20 | 74 | 475 | 54.00 | 154.82 | Rieske 2Fe-2S domain-containing protein [*Micrococcales*] | WP_228164524 | 100.00% |
| 3004 (ACQVDU_13245) | 2725684-2726598 | 15 | 63 | 304 | 32.10 | 129.57 | DMT family transporter [*Microbacterium*] | WP_228178617 | 93.09% |
| 2349 | 2025295-2026383 | 13 | 45 | 362 | 37.60 | 111.50 | threonine synthase | WP_300594141 | 100.00% |

| | | | | | | | | | |
|---|---|---|---|---|---|---|---|---|---|
| (ACQVDU_09975) | | | | | | | [*Microbacterium*] | | |
| 2584 (ACQVDU_11170) | 2277460-2278902 | 7 | 32 | 480 | 52.90 | 89.93 | cytochrome ubiquinol oxidase subunit I [*Microbacterium*] | WP_300591869 | 99.79% |
| 803 (ACQVDU_04060) | 823821-824276 | 8 | 30 | 151 | 15.90 | 80.38 | type II 3-dehydroquinate dehydratase [*Microbacterium*] | WP_300592868 | 100.00% |
| 1724 (ACQVDU_06920) | 1393943-1395784 | 7 | 34 | 613 | 68.30 | 67.39 | PEP-utilizing enzyme [*Microbacterium*] | WP_336401120 | 88.89% |
| 1704 (ACQVDU_06830) | 1378368-1379459 | 9 | 27 | 363 | 37.9 | 54.91 | glycerol dehydrogenase [*Herbiconiux*] | WP_092551598 | 58.47% |
| 2521 (ACQVDU_10845) | 2213638-2215359 | 10 | 29 | 573 | 60.40 | 43.57 | acyl-CoA dehydrogenase [*Microbacterium*] | WP_331791808 | 98.60% |
| 1242 (ACQVDU_15265) | 3100608-3102155 | 5 | 34 | 515 | 54.80 | 37.58 | alpha/beta hydrolase [*Microbacterium*] | WP_331791728 | 99.61% |

| | | | | | | | | | |
|---|---|---|---|---|---|---|---|---|---|
| 2441 (ACQVDU_10450) | 2126855-2128228 | 10 | 22 | 457 | 50.4 | 29.95 | FAD-dependent oxidoreductase [*Microbacterium*] | WP_292710698 | 100.00% |
| 308 (ACQVDU_01550) | 315433-316728 | 7 | 24 | 431 | 48.00 | 26.10 | citrate synthase [*Microbacterium*] | WP_363497997 | 99.54% |
| 2404 (ACQVDU_10260) | 2089875-2090711 | 7 | 26 | 278 | 28.40 | 21.48 | pyrroline-5-carboxylate reductase [*Microbacterium*] | WP_300591385 | 98.56% |
| 2612 (ACQVDU_11315) | 2307237-2308601 | 8 | 20 | 454 | 46.8 | 15.93 | 3-phosphoshikimate 1-carboxyvinyltransferase [*Microbacterium*] | WP_363495956 | 97.36% |

a. Number of different peptides identified. The higher the number, the higher the protein abundance.

b. Number of peptides matched to secondary spectra.

c. Number of amino acids.

d. Protein matching score, the higher the score, the higher the confidence

e. The top BLASTP hit was selected from NCBI Non-redundant Protein Sequences Database.
